# Disparate evolution of virus populations in upper and lower airways of mechanically ventilated patients

**DOI:** 10.1101/509901

**Authors:** Björn F. Koel, Frank van Someren Gréve, René M. Vigeveno, Maarten Pater, Colin A. Russell, Menno D. de Jong

## Abstract

In routine surveillance and diagnostic testing, influenza virus samples are typically collected only from the upper respiratory tract (URT) due to the invasiveness of sample collection from the lower airways. Very little is known about virus variation in the lower respiratory tract (LRT) and it remains unclear if the virus populations at different sites of the human airways may develop to have divergent genetic signatures. We used deep sequencing of serially obtained matched nasopharyngeal swabs and endotracheal aspirates from four mechanically ventilated patients with influenza A/H3N2 infections. A physical barrier separating both compartments of the respiratory tract introduced as part of the medical procedures enabled us to track and compare the genetic composition of the virus populations during isolated evolution in the same host. Amino acid variants reaching majority proportions emerged during the course of infection in both nasopharyngeal swabs and endotracheal aspirates, and amino acid variation was observed in all influenza virus proteins. Genetic variation of the virus populations differed between the URT and LRT and variants were frequently uniquely present in either URT or LRT virus populations of a patient. These observations indicate that virus populations in spatially distinct parts of the human airways may follow different evolutionary trajectories. Selectively sampling from the URT may therefore fail to detect potentially important emerging variants.

**Importance:** Influenza viruses are rapidly mutating pathogens that easily adapt to changing environments. Although advances in sequencing technology make it possible to identify virus variants at very low proportions of the within-host virus population, several aspects of intrahost viral evolution have not been studied because sequentially collected samples and samples from the lower respiratory tract are not routinely obtained for influenza surveillance or clinical diagnostic purposes. Importantly, how virus populations evolve in different parts of the human respiratory tract remains unknown. Here we used serial clinical specimens collected from mechanically ventilated influenza patients to compare how virus populations develop in the upper and lower respiratory tract. We show that virus populations in the upper and lower respiratory tract may evolve along distinct evolutionary pathways, and that current sampling and surveillance regimens likely capture only part of the complete intrahost virus variation.

## Introduction

Influenza A viruses are genetically variable due to the low fidelity of the influenza virus RNA polymerase and can therefore quickly evolve in response to changing environments and selection pressures, leading to escape from natural or vaccine-induced immunity, antiviral drug resistance and adaptation to new hosts. Intrahost influenza virus populations are genetically heterogeneous and consist of closely related but diverse virus variants. The evolution of influenza viruses has been studied in much detail at population levels on a global scale, but the intrahost evolutionary processes that are fundamental to emergence of new variants in the human population are less well understood.

Influenza viruses can infect and replicate in epithelial cells throughout the upper (URT) and lower respiratory tract (LRT) (1, 2). Differences between cell types and receptor distribution along the respiratory tract, as well as local conditions such as temperature and immunity, may favor the emergence of different variants in different compartments of the human airways. Recent studies indeed suggest compartmentalization of seasonal influenza viruses in the URT and LRT. Richard *et al.* reported that intranasal and intratracheal co-inoculation of ferrets resulted in minimal reassortment of the inoculated viruses (3), and Yan *et al.* found that URT and LRT infections appeared to behave as independent virus populations based on aerosol shedding of human volunteers (4). Although these studies provide clues that virus populations in spatially separated compartments of the respiratory tract may evolve independently, current insights into potential differences in genetic composition and evolution of virus populations between URT and LRT compartments remain incomplete.

Improved understanding would ideally require analyses of serial specimens collected in parallel from both respiratory compartments during the course of influenza, the feasibility of which is challenging due to the invasiveness of such sampling. We had an opportunity to perform such analyses in four influenza A/H3N2 virus-infected patients who participated in a study of the prevalence and shedding patterns of respiratory viruses in critically ill patients receiving mechanical ventilation (5, 6). For the purpose of this study, nasopharyngeal specimens and endotracheal aspirates were collected each day from patients while intubated and ventilated. Endotracheal tubes used for mechanical ventilation contain a balloon cuff designed to provide a seal inside the trachea that allows airflow through the tube whilst preventing passage of air or fluids around it. Importantly, this feature allowed analysis of virus populations in upper and lower respiratory tract compartments in isolation. Using a next generation sequencing approach, virus populations from both compartments were thus characterized over time, providing evidence indicating differences in genetic composition and diversity.

## Results

### Limited genetic variation in virus populations from nasopharyngeal swabs and endotracheal aspirates

Matched nasopharyngeal swabs (NPS) and endotracheal aspirates (ETA) were collected daily from mechanically ventilated patients in the context of a Dutch multicenter observational clinical study investigating the prevalence and shedding patterns of respiratory viruses in critically ill patients (5, 6). For the current study we selected all patients with laboratory-confirmed influenza A/H3N2 virus infections in whom viral RNA could be detected in paired NPS and ETA specimens during at least 2 consecutive days. In NPS and ETA from the four patients thus included, influenza virus RNA was detected during respectively 3 - 7 days and 3 - 26 days following intubation (Fig. 1, Fig. S1). Whole genome next generation sequencing (NGS) was performed directly on all virus positive samples to quantify the within-host genetic variation and changes in amino acid variant frequency over time. Genomic data of sufficient quality were obtained from 51 samples in total. To limit inclusion of artifactual variants introduced during library preparation or sequencing, variants were only considered for analyses if present in the virus population during multiple days.

**FIG 1.**
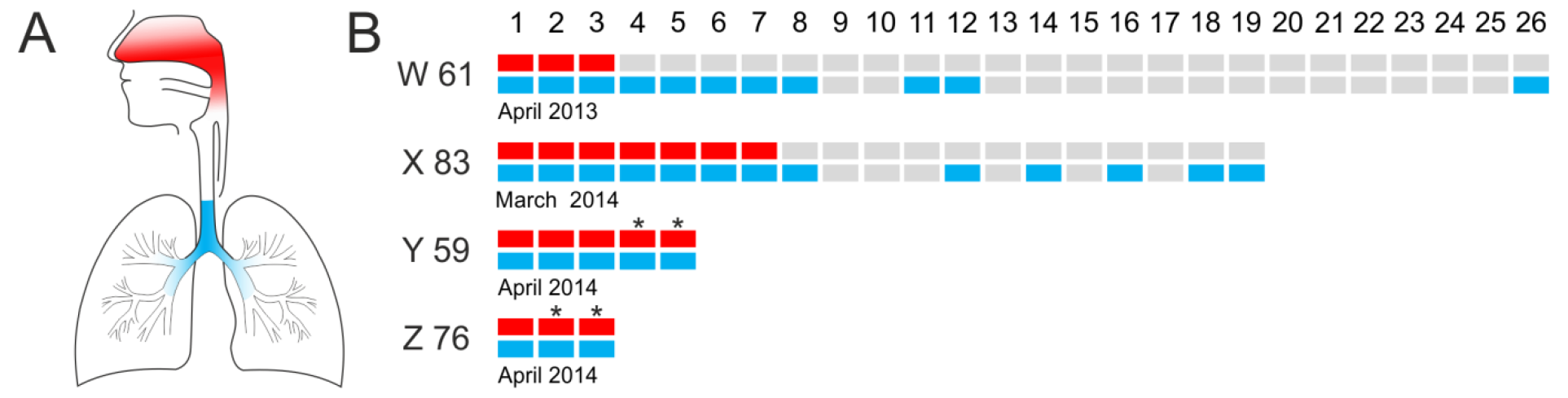
Sample overview (A) Samples were collected daily from the nasopharyngeal (red) and bronchotracheal area (blue) and are referred to here as upper and lower respiratory tract samples, respectively. (B) High quality NGS data were obtained from serially collected samples throughout the course of infection of four patients; W, X, Y, and Z. Samples indicated as grey bars were influenza virus negative or NGS data did not pass quality control during analysis. Numbers in the top row of panel B indicate days since intubation. Patient designation, age and admission date are indicated in the figure. An asterisk indicates the days on which antiviral treatment with oseltamivir was administered.

There was limited but evident genetic variation in the intrahost virus populations in the four patients. We identified 31 amino acid variants in the NPS and 34 amino acid variants in the ETA collected among the four patients (Fig 2). Variations in NPS occurred on 28 unique amino acid positions on all but the M1 and NP proteins (Fig 2A). In ETA, variants on 26 unique amino acid positions were found and occurred on all but the PA-X, NS2, and PB1-F2 proteins (Fig. 2B). To identify temporal differences in variant proportions, we arbitrarily set a 5% threshold value for what constitutes a substantial variant proportion difference between same-patient NPS or ETA samples. The difference between minimum and maximum variant proportions exceeded this threshold for 20 variants in NPS and 28 variants in the ETA (Figs. 2C and 2D, Figs S2–S5). Changes in variant proportion fluctuated over time. Variant proportion differences of 15 - 20% between consecutive days were observed in all four patients (Figs S2–S5). However, most variants showed short-lived changes in variant proportion, typically reaching 10 - 20% in the virus population. Few variants persisted at low proportions (<10%) for several days or reached proportions above 20%. Four variants reached majority in the virus population (>50%)—HA G5E (NPS, patient X), NS1 H59L (NPS, patient X), NA E76D (NPS, patient Y), and NS1 L185F (ETA, patient W). However, consistent outgrowth during sampling was observed only for NS1 H59L, whereas the variant proportions of HA G5E and NS1 L185F dropped below the detection limit within days after reaching majority. We could not determine evolutionary patterns for the duration of infection for NA E76D because the coverage of this genetic region was below our inclusion threshold for three out of five days.

**FIG 2.**
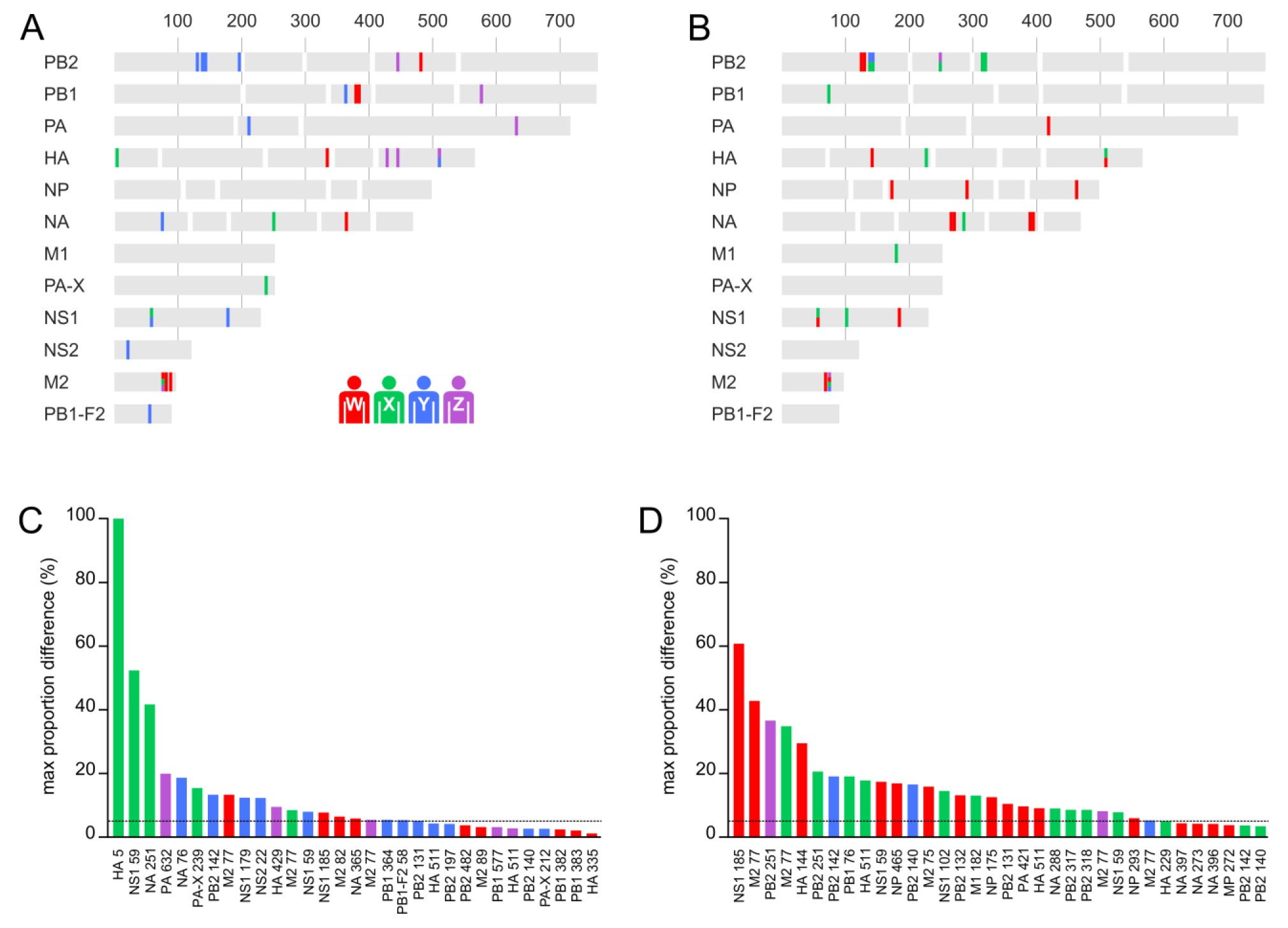
Overview of proteins and amino acid positions responsible for diversity in the virus populations in NPS and ETA. Variants are color coded by patient, patient W; red, patient X; green, patient Y; blue, patient Z; purple. (A) Variable amino acid positions identified in NPS indicated on the A/H3N2 virus proteins. Vertical lines indicate amino acid positions at intervals of 100 amino acids. White vertical bars indicate primer regions. (B) As panel A, but showing the variable amino acid positions identified in ETA. (C) Maximum proportion differences of variants in NPS. X-axis labels indicate protein and amino acid positions. The threshold used in the analysis of temporal variation is indicated as a dashed line. Variants with temporal differences in variant proportion exceeding this threshold are listed in Table 1. (D) As panel B, but showing maximum proportion differences of variants in ETA. A/H3N2 virus HA numbering according to Burke *et al.* (28).

**Table 1.** Amino acid variants with temporal differences in variant proportion exceeding 5%.

| patient | sample type |  |  |  |  |  |
| --- | --- | --- | --- | --- | --- | --- |
|  | swabs |  |  | aspirates |  |  |
| W | PB2 482KR | HA I335K | M2 R77stop | PB2 G131V | NP H272C | NA I397Q |
|  | PB1 N382H | NA T365I | M2 N82S | PB2 P132L | NP 273KR | NS1 H59L |
|  | PB1 E383V | NS1 L185F | M2 S89G | PA V421I | NP K293Q | NS1 L185F |
|  |  |  |  | HA N144K | NP E465V | M2 E75R |
|  |  |  |  | HA Y511D | NA V396A | M2 R77stop |
|  |  |  |  | NP R175K |  |  |
| X | HA G5E | NS1 H59L |  | PB2 K140I | PB1 D76N | NS1 H59L |
|  | NA V251I | M2 R77stop |  | PB2 R142F | HA R229K | NS1 W102C |
|  | PA-X T239M |  |  | PB2 R251K | HA Y511D | M2 R77stop |
|  |  |  |  | PB2 L317F | NA R288L |  |
|  |  |  |  | PB2 R318S | M1 A182T |  |
| Y | PB2 G131V | HA Y511D | NS2 G22R | PB2 K140I |  |  |
|  | PB2 K140I | NA E76D | PB1-F2 W58stop | PB2 R142F |  |  |
|  | PB2 R142F | PA-X V212A |  | M2 R77stop |  |  |
|  | PB2 K197N | NS1 H59L |  |  |  |  |
|  | PB1 L364I | NS1 G179E |  |  |  |  |
| Z | PB1 K577N | HA V429I | M2 R77stop | PB2 R251K |  |  |
|  | PA S632T | HA Y511D |  | M2 R77stop |  |  |

The influenza virus proteins analyzed for this study are basic polymerase 2 (PB2), basic polymerase 1 (PB1), acidic polymerase (PA), hemagglutinin (HA), nucleoprotein (NP), neuraminidase (NA), matrix protein 1 (M1), acidic polymerase protein X (PA-X), non-structural protein 1 and 2 (NS1, NS2), matrix protein 2 (M2), and basic polymerase 1 frame 2 (PB1-F2). A/H3N2 virus HA numbering according to Burke *et al.* (28).

### Virus populations in nasopharyngeal swabs and endotracheal aspirates are genetically diverse

To determine whether the genetic composition of the virus populations in NPS and ETA was different, we next analyzed variant proportions in time- and patient matched NPS and ETA. For this analysis we included the variants that were detectable during multiple days and where the difference between variant proportions in NPS and ETA exceeded 5%.

Panels A and B of figure 3 show for each of the patients the variants present in NPS and ETA at the first day of sample collection. Variants that met our inclusion criteria and that were unique to either NPS or ETA were observed in all four patients, albeit at moderate proportions. Variants present in both NPS and ETA similarly showed moderate differences in variant proportions between NPS and ETA samples. Because our analysis of temporal differences in variant proportion indicated that the virus populations in these patients evolved over time, we repeated the analysis using for each patient the samples collected at the last time point where influenza virus-positive NPS and ETA were available (Fig. 3 panels C and D). Also here, variant proportions in the NPS and ETA were genetically different in all patients, and included variants that were present in only one airway compartment. These results show that the genetic composition of the virus populations in NPS and ETA was different during identical periods after intubation in all four patients.

**FIG 3.**
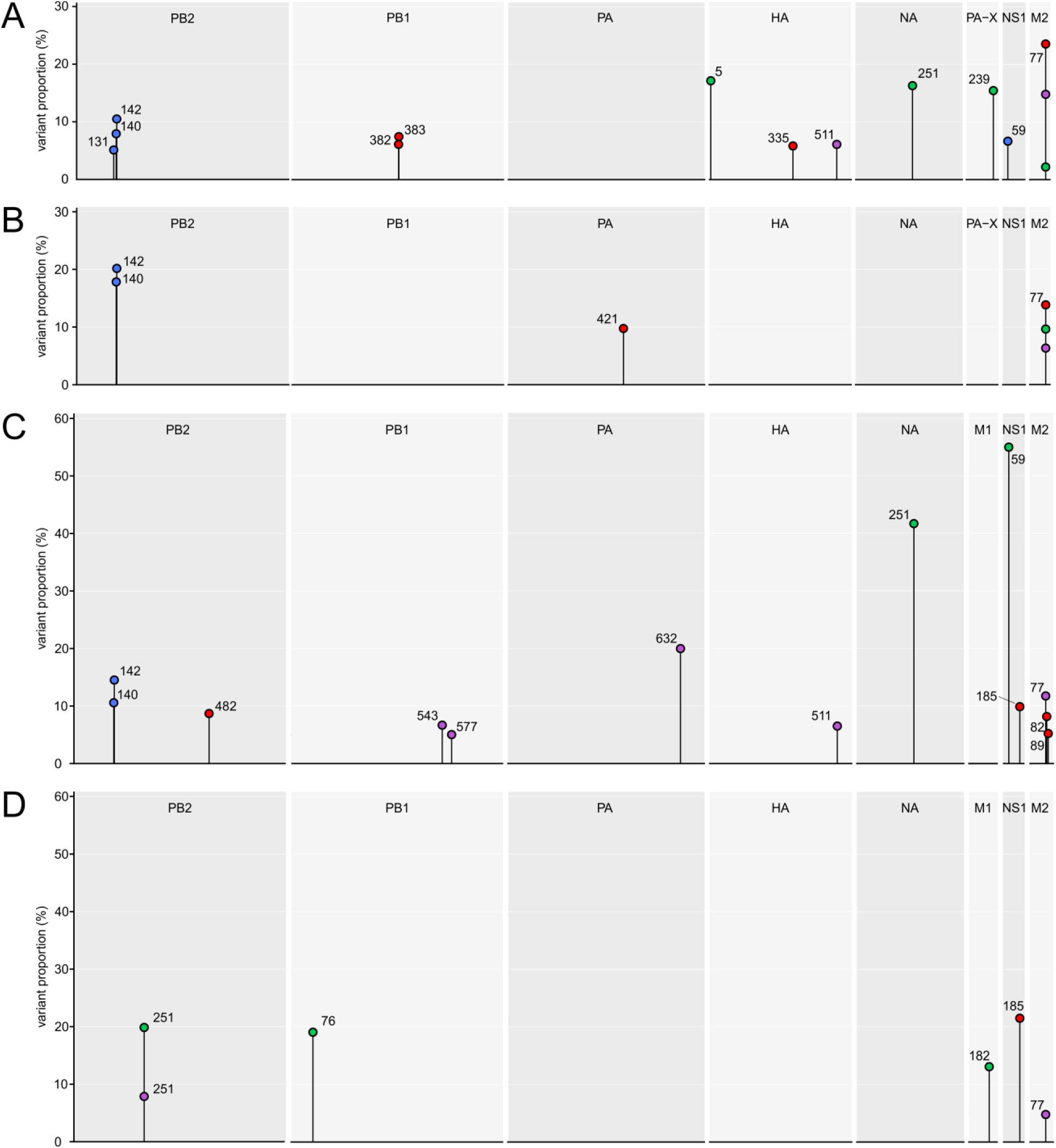
Variant proportions in NPS and ETA at matched time points. Variants are color coded by patient, patient W; red, patient X; green, patient Y; blue, patient Z; purple. Panels (A) and (B) show the variant proportions in NPS and ETA, respectively, at the first day of sample collection. Column headings indicate the proteins in which variation was detected. Variants are labelled by amino acid position. Panels (C) and (D) show the variant proportions of variants in NPS and ETA, respectively, from the last available influenza virus-positive time-matched samples.

The analysis of differences between the virus populations in NPS and ETA revealed a number of notable variants. Two of the four variants that reach majority proportions, HA G5E and NA E76D, were uniquely observed in the NPS of patients X and Y, respectively (Figs. S3 and S4). The remaining two variants that reach majority proportions, NS1 H59L and NS1 L185F, reach peak variant proportions while being undetectable in the other airway compartment at matched time points. In addition, these variants were present and continued to vary in proportion in ETA after the virus became undetectable in NPS, as was also the case for HA N144K (Fig. 4, and Figs. S2 and S3). Finally, a variant with a truncation of the M2 cytoplasmic tail, M2 R77stop, was observed in all four patients and was absent only from NPS from patient Y (Figs. S2–S5). Variant proportions of M2 R77stop were mostly different between same-patient time-matched NPS and ETA samples. Like the variants outlined in Fig 4, M2 R77Stop remained present and reached high variant proportions in ETA samples of patients W and X after NPS samples became virus negative.

**FIG 4.**
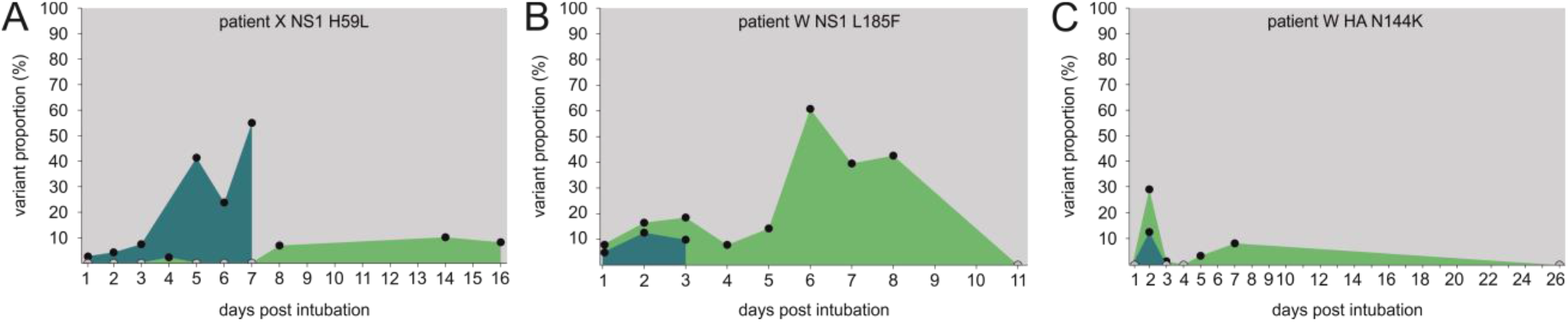
Sustained evolution of variants in the LRT. Blue and green filled areas indicate the proportion of the variant in the URT and LRT, respectively. Filled circles represent the samples included in variant analysis; black dots represent samples with variant proportions above the detection limit, grey dots indicate absence of the variant. (A) evolutionary dynamics of NS1 H59L in patient X. (B) evolutionary dynamics of NS1 L185F in patient W. (C) evolutionary dynamics of HA N144K in patient W.

## Discussion

Meaningfully studying intrahost genetic variation of human influenza A virus populations has become possible in the past few years as a result of progress in sequencing technologies. Multiple groups have now used deep sequencing approaches to determine the genetic variation of human influenza A viruses in patient-derived materials. These studies have mostly focused on variation in relation to immune escape because of the major role of this process in influenza virus evolution (7–9), or on better understanding transmission bottlenecks and stringency of intrahost selection pressures (10–12). The scarcity of sequentially collected clinical specimens accessible for scientific purposes, particularly those obtained from the lower respiratory tract, has limited the opportunities for studying the temporal genetic variation of within-host influenza virus populations (9, 13, 14). Here we had the opportunity to study intrahost viral genetic variation in longitudinally collected nasopharyngeal swabs and endotracheal aspirates of individuals in whom upper and lower airway compartments were separated by a physical barrier due to mechanical ventilation. We evaluated the intrahost evolutionary patterns of influenza A/H3N2 virus in both airway compartments and compared the genetic composition of the virus populations in the upper and lower airways.

Our results indicate intrahost viral genetic diversity and temporal differences in the composition of the virus populations in the URT and in the LRT of all patients. The modest number of variants observed in each patient is in concordance with previous studies that reported limited genetic variation based on studies using single time points (7, 8, 10). Nonetheless, we observed outgrowth of several variants towards or above majority proportions, often reaching peak proportions within a week after the start of sample collection. Our data also point out that variants may reach majority proportions in both airway compartments. These outcomes suggest that substantial genetic changes in the influenza A virus populations of the URT and LRT can take place during the short time periods typical of transient influenza virus infections.

The different variants and variant proportions detected in URT and LRT samples indicates that genetically related but distinct virus populations existed in the upper and lower airway compartments of the patients included in this study. Our finding that distinct viral genotypes frequently existed in one but not the other airway compartment complements a recent study that demonstrated spatial separation of the virus populations in humans without the mechanical barrier that was present in the patients sampled for the current study (4). That work showed that humans generate infectious aerosols that represent infection of the LRT and that viral load in URT samples poorly predicted shedding of virus in aerosols. Here we show that virus populations in distal parts of the human airways were composed of distinct viral genotypes that followed isolated evolutionary pathways. If aerosols support influenza virus transmission, the results of the current study imply that variants originating from the lower airways may contribute to long-term (interhost) virus evolution. The differences between genetic compositions of virus populations in the URT and LRT, even during matched time points, suggests that genetic analysis of the virus population in URT samples is a poor proxy for genetic variation in the LRT virus population. Given that current sampling regimens rarely include sample collection from the LRT, a potentially important evolutionary setting may be almost entirely unexplored.

While it is plausible that the distinct physiological conditions and receptor distributions in the URT and LRT gives rise to site specific variants, identification of adaptive amino acid changes resulting from selection pressures specific to either compartment was precluded by the limited number of patients in this study. However, HA N144K, HA R229K, NA V251I, and NS1 L185F have been associated with antibody evasion or immune modulation (15–20), which may explain the rapid increase in variant proportion of some of these variants. Our findings are in agreement with previous results that reported limited variation on amino acid positions associated with immune escape (7–9, 12), and studies that suggested that antigenic adaptation is unlikely to be a major mechanism responsible for intrahost genetic variation (8). Additionally, truncation of M2 at position 77 was shown to have no impact on ion-channel activity (21), suggesting that this amino acid variation may also have been phenotypically neutral in the viruses observed in this study. The phenotypic effects of other notable variants, including those reaching majority proportions, have to our knowledge not been reported.

In summary, we showed disparate evolution of virus populations in spatially separated parts of the human airways following natural infection with a seasonal human influenza virus. The temporal variability and genetic differences between intrahost virus populations puts into question the significance of samples collected at a single site or at a single time point. Timing, duration, and site of sample collection are critical variables that could affect the outcomes of influenza surveillance, and of studies into influenza disease and virus evolution. This is especially true for studies looking into intrahost genetic diversity using deep sequencing approaches.

## Materials and Methods

### Patients and samples

The samples used in this study were collected as part of a multicenter prospective observational study performed in The Netherlands (Dutch Trial Register NTR4102) as described previously (22). A waiver from the Medical Research Involving Humans Act was provided by the Institutional Review Board of the Academic Medical Center, Amsterdam, due to non-invasiveness of study procedures. Patients and/or their legal representatives were provided with written study information at ICU admission, and could opt-out of the study participation. Included were critically ill patients requiring intubation and mechanical ventilation, admitted to the participating ICUs between April 2013 and April 2014. Daily nasopharyngeal NPS and tracheobronchial ETA were collected until detubation or death while on mechanical ventilation. All samples obtained upon admission were tested with a validated multiplex RT-PCR for respiratory viruses, as previously described (23, 24). For the current study, all patients that were influenza A/H3N2 virus positive for multiple days positive were included. Intubation and sampling of patients X,Y, and Z started at the day of admission to the ICU. Patient W had been admitted to the ICU for 6 days prior to the start of intubation and sampling. Patient characteristics are available from Table S1. Viral loads for all patients and time points are indicated in figure S1.

### Library preparation and deep sequencing

Total RNA was extracted from the clinical specimens using the High Pure RNA isolation kit (Roche, 11828665001) according to manufacturer’s instructions. Influenza RNAs were reverse transcribed and amplified using the superscript III One-Step RT PCR Platinum Taq High Fidelity DNA Polymerase (ThermoFisher, 12574030) and A(H3N2) virus subtype and gene segment specific primers (Table S3). For whole genome amplification we performed 20 independent PCR reactions in total. Three partly overlapping amplicons were generated for the PB2, PB1, PA, HA, NA and NP segments each, a single amplicon each was generated for the M and NS gene segments. For each sample, PCR products were pooled in equimolar concentrations and subsequently purified using Agencourt Ampure XP beads (Beckman Coulter, A63882) and quantified using the Qubit dsDNA HS assay kit (ThermoFisher, Q32854). Pooled and cleaned amplicons were diluted to 0.2 ng/μl for subsequent library preparation.

Sequencing libraries were prepared using the Nextera XT DNA Library Preparation kit (Illumina, FC-131-1096) according to manufacturer’s instructions. Briefly, for each sample 5 μl of diluted amplicons were enzymatically fragmented and Illumina adapters were ligated to the fragments. Subsequently each sample was purified twice using Agencourt Ampure XP beads. Library size distribution was evaluated using the High sensitivity dsDNA kit on a 2100 Bioanalyzer (Agilent, 5067-4626) and qPCR based library quantification was performed using the KAPA Library Quantification kit for Illumina platforms (KAPA Biosystems, KK4824) on a LightCycler480 (Roche). Normalized library pools were sequenced on an Illumina MiSeq machine using the 600-cycle MiSeq Reagent Kit v3 (Illumina, MS-102-3003). All FASTQ files are available on request.

### Quality control, variant detection and data analysis

Quality trimming of Illumina MiSeq reads was performed using the Maximum Information quality filtering approach of the Trimmomatic tool (version 0.36, parameters; leading:3, trailing:3, maxinfo: 80:0.4, crop:280) (25). Merging, mapping, and coverage analysis was done using the BBmerge, BBwrap, and pileup scripts from the BBMap bioinformatics toolkit version 36.27 (26). Read pairs with inappropriate orientation and reads with a Q-score below 25 were discarded. Quality control was monitored using FastQC version v0.11.5 (27).

Subsequent steps were performed using a set of custom scripts (available on request). Mapped reads were translated and prepared for variant calling by identification of appropriate reading frames and conversion of read numbering to protein numbering for the 12 influenza A virus proteins considered here; PB2 (polymerase basic 2), PB1 (polymerase basic 1), PA (polymerase acidic), HA (hemagglutinin), NP (nucleoprotein), NA (neuraminidase), M1 (matrix protein 1), PAX (polymerase acidic protein-X), NS1 (non-structural protein 1), NS2 (nuclear export protein), M2 (matrix 2 ion channel), and PB1-F2 (PB1 frame 2). HA amino acid positions in the manuscript are numbered according to Burke *et al.* (28). For each patient the data on coverage per position, variant count, and variant proportion was collected for all time points and for URT and LRT samples. Next a number of filtering steps was performed. Variants passing filter were outside of primer regions, had a minimum coverage of 100x for each position in the codon, and were present at at least 1% of the total virus population with a minimum of five observations per sample. All variants included in our analyses were detectable in two or more samples from the same airway compartment and reached a variant proportion of at least 5%. Variants identified in the overlapping regions of the PA gene products (PA and PA-X), M gene products (M1 and M2), and NS gene products (NS1 and NS2) were called independently.

## Acknowledgements

This research was supported by ZonMW TOP grant and a Postdoc Stipend of the Amsterdam Infection and Immunity Institute. The authors are grateful to Sylvie Koekkoek, Silvana Roos, Matthijs Welkers, Dirk Eggink, Nicole Juffermans, and Marcus Schultz from the AMC Amsterdam, The Netherlands, and to Frank Harders from the Central Veterinary Institute, The Netherlands, for assistance and technical support. BFK and MP thank SURFsara (www.surfsara.nl) for the support in using the Lisa Compute Cluster.

---

**FIG S1.**
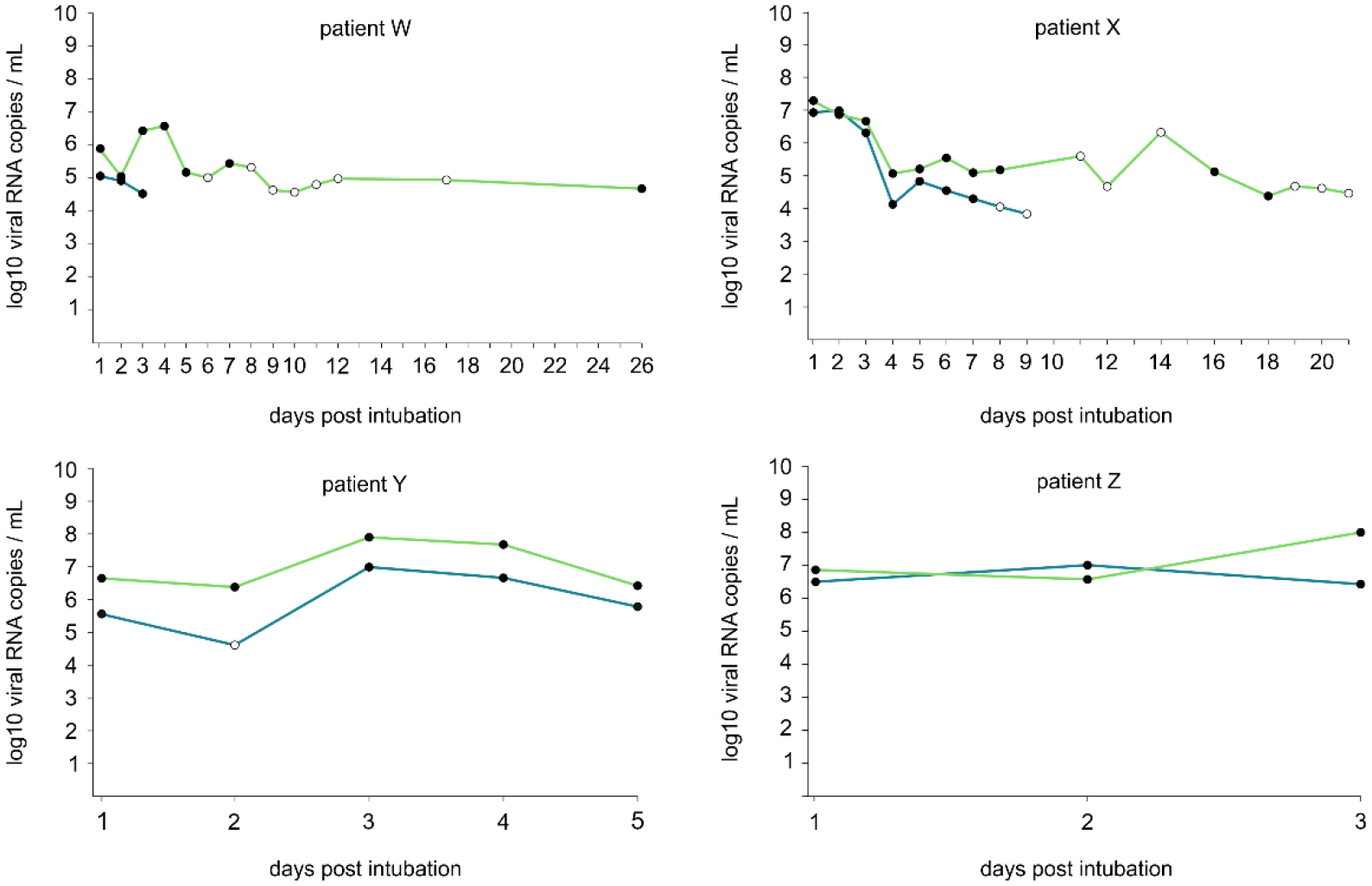
Viral load in NPS (blue) and ETA (green) specimens of patients W, X, Y, and Z. Filled circles indicate high quality samples included in variant analysis.

**FIG S2.**
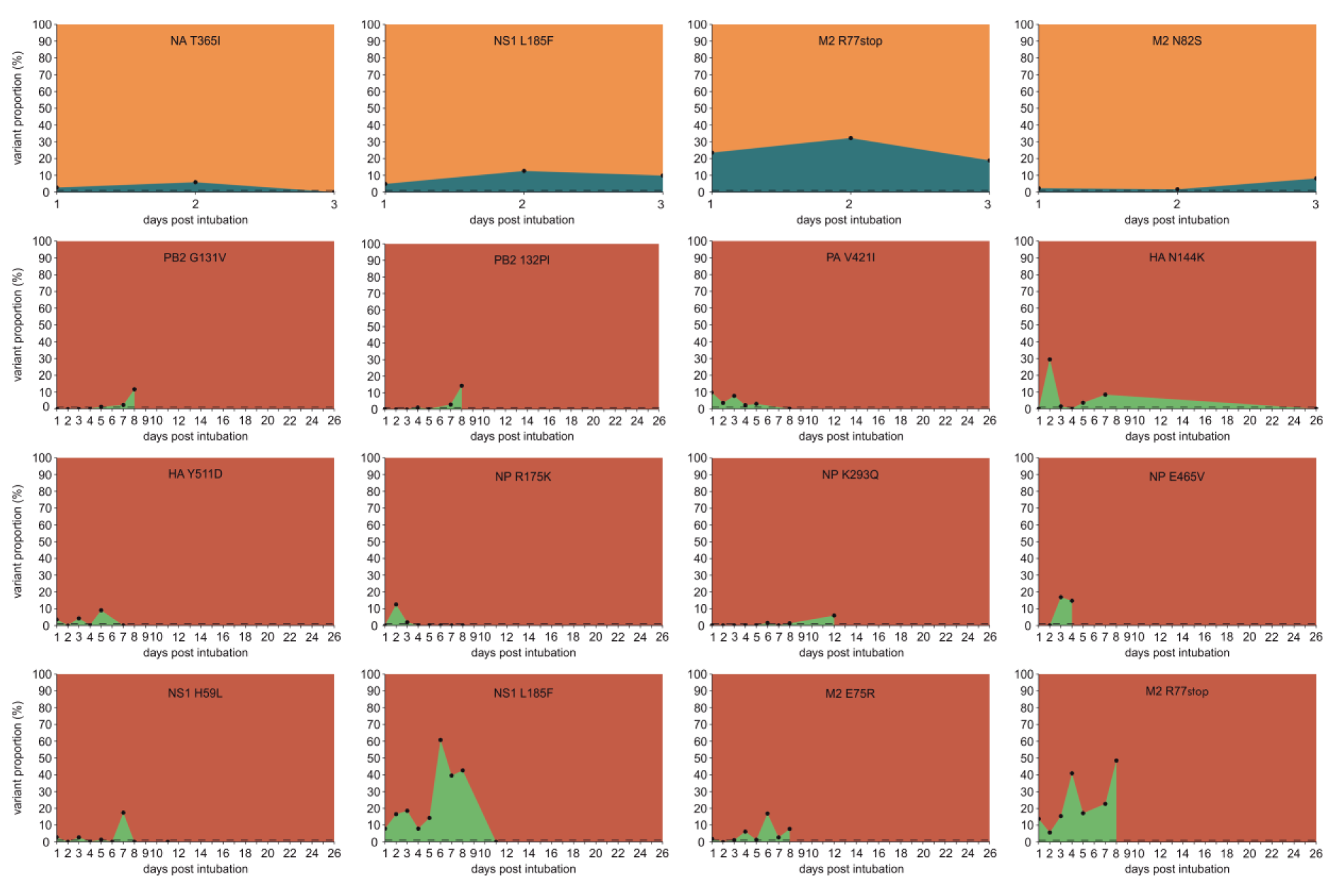
Evolutionary dynamics of variants showing substantial temporal variation for patient W. Blue on an orange background indicates the proportion of the variant in the URT, green on a brown background indicates the proportion of the variant in the LRT. Filled circles represent the samples included in variant analysis. The protein and variant are indicated in the figures. Capital letters indicate the amino acid that is the majority amino acid on a given position, small letters indicate the minority variant.

**FIG S3.**
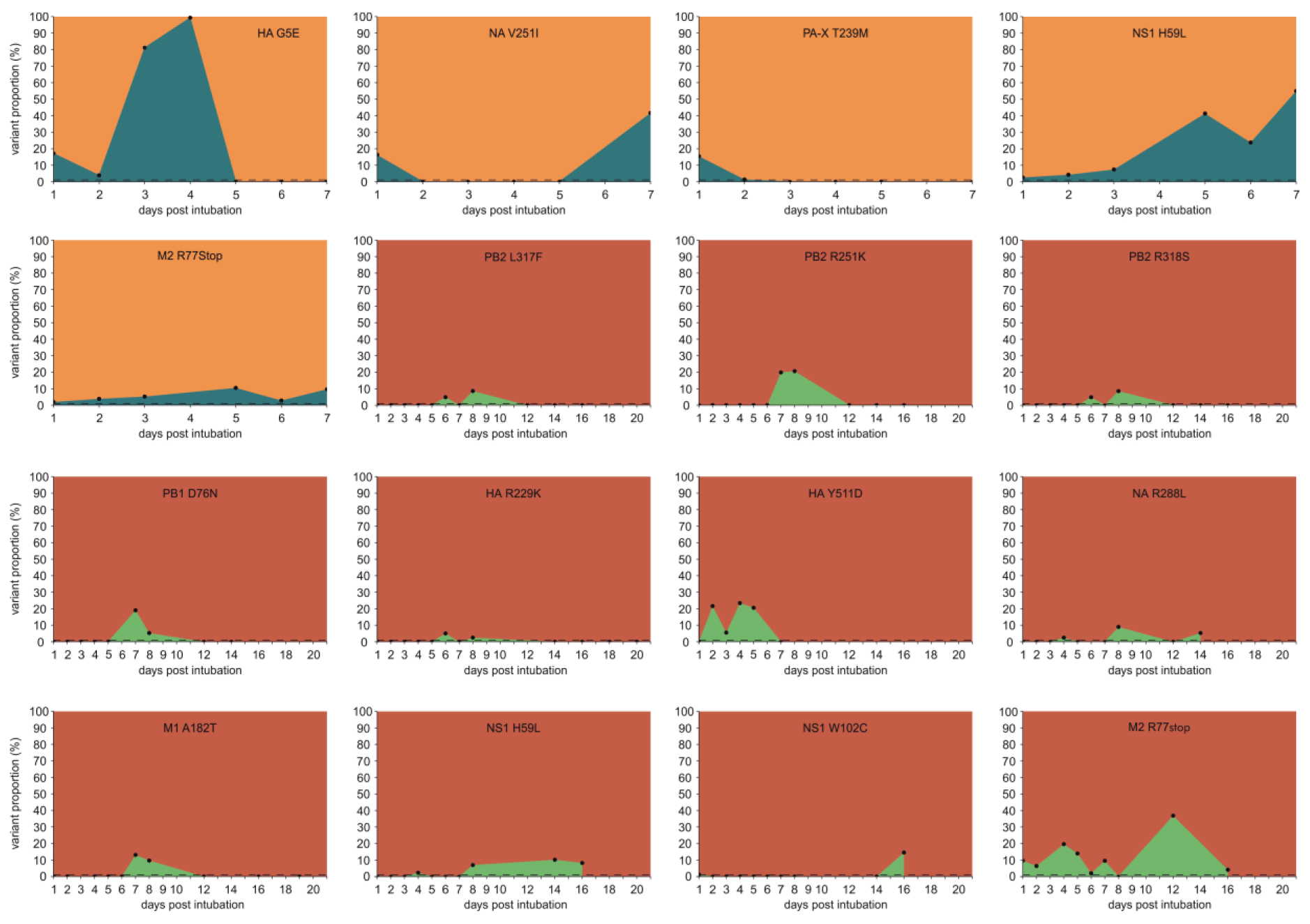
Evolutionary dynamics of variants showing substantial temporal variation for patient X. Symbols and colors as in Fig S2.

**FIG S4.**
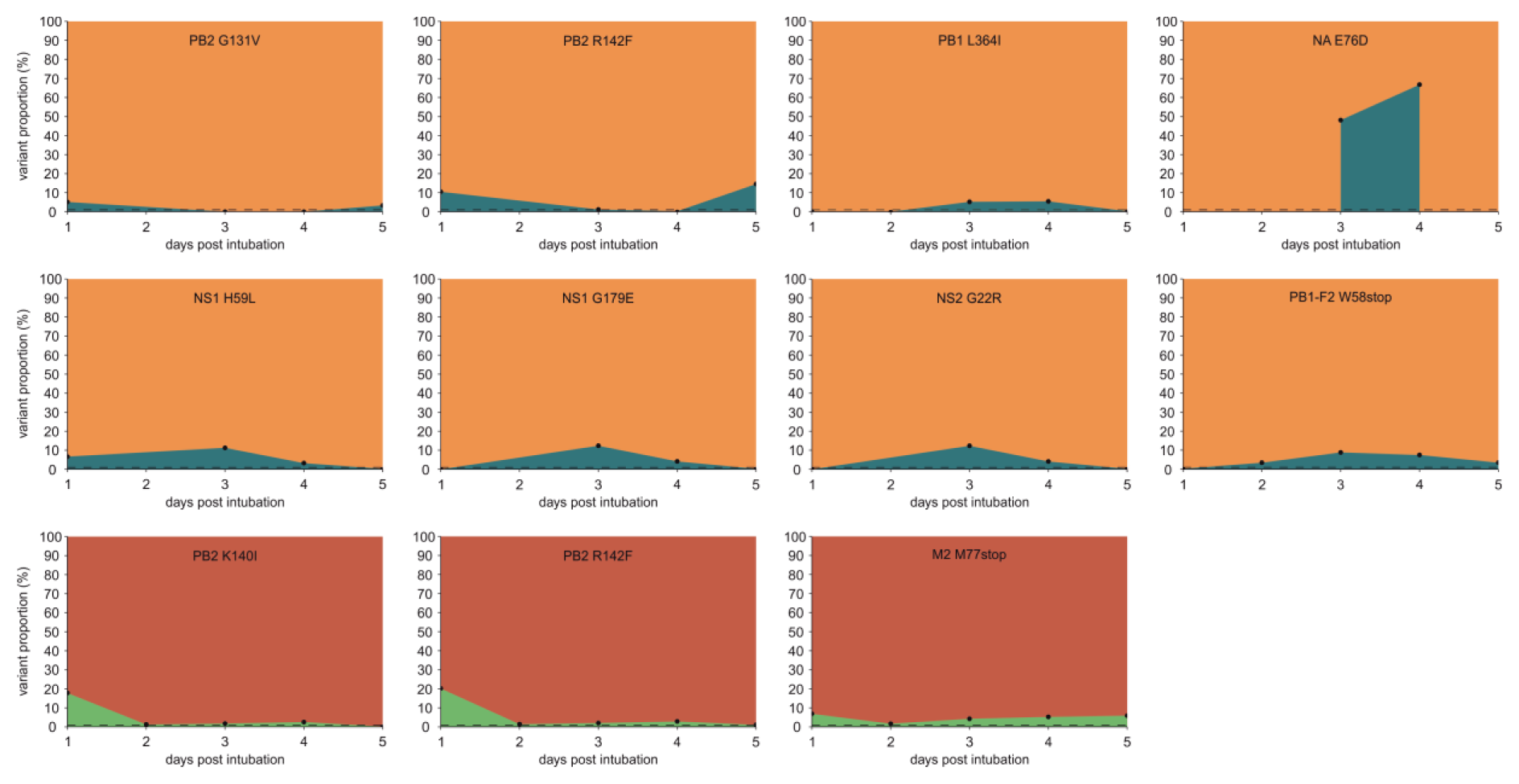
Evolutionary dynamics of variants showing substantial temporal variation for patient Y. Symbols and colors as in Fig S2.

**FIG S5.**
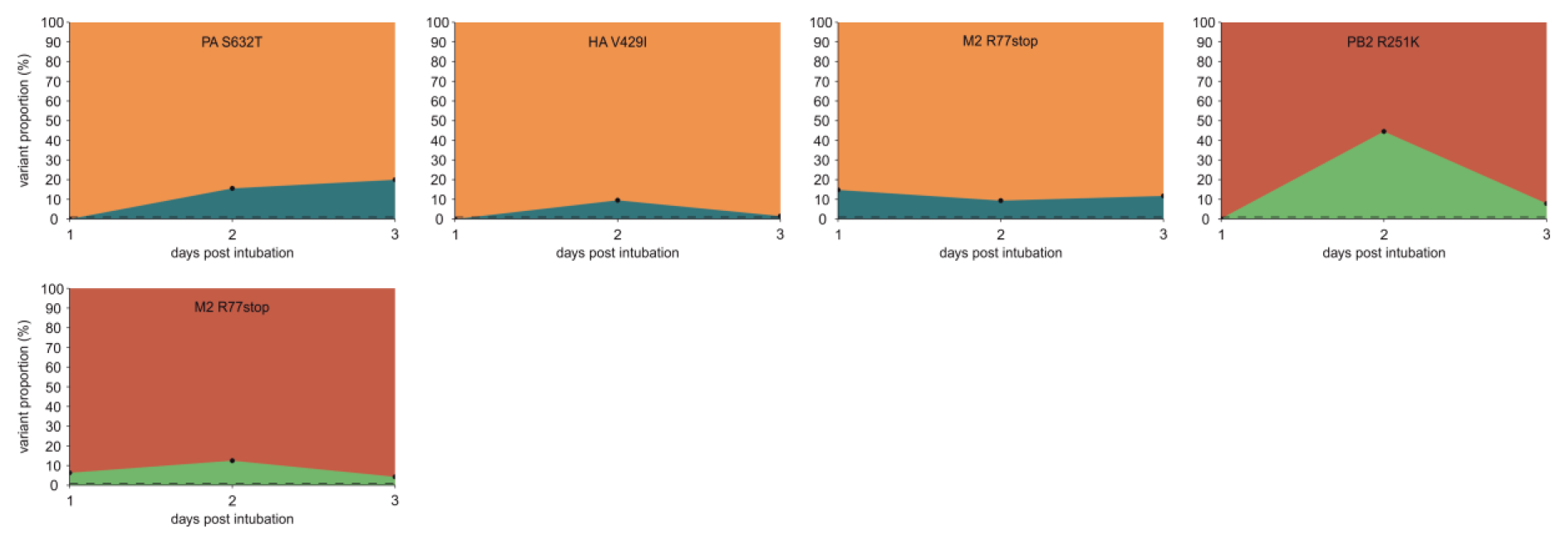
Evolutionary dynamics of variants showing substantial temporal variation for patient Z. Symbols and colors as in Fig S2.

**Table S1.** Patient characteristics.

|  | patient W | patient X | patient Y | patient Z |
| --- | --- | --- | --- | --- |
| Reason of ICU admission | Severe acute respiratory infection | Severe acute respiratory infection | Respiratory surgery | Severe acute respiratory infection |
| Comorbidities | none | none | none | DMII, CHF |
| Age (years) | 61 | 83 | 59 | 76 |
| Sex | male | male | female | female |
| Received oseltamivir | no | no | yes | yes |
| Clinical parameters (at ICU admission) |  |  |  |  |
| Temperature (°C) | 38.9 | 36.5 | 36.7 | 34.3 |
| Leucocytes (10<sup>9</sup>/L) | 10,6 | 12,4 | 11,9 | 24,8 |
| CRP (mg/L) | 11 | 37 | 11 | 41 |
| Consolidation on chest X-ray | no | no | no | yes |
| APACHE II score | 24 | 30 | 14 | 35 |
| SAPS II | 50 | 55 | 39 | 62 |
| Clinical outcomes |  |  |  |  |
| ICU length of stay (days) | 37 | 26 | 6 | 4 |
| Hospital length of stay (days) | 86 | 34 | 13 | 4 |
| Died during hospital admission | no | yes | no | yes |

Abbreviations: APACHE II = Acute Physiology and Chronic Health Evaluation II ; CHF = congestive heart failure; CRP = C-reactive protein; DMII = diabetes mellitus type II; ICU = intensive care unit; SAPS II = Simplified Acute Physiology Score II;

